## Supplementary material for "RST1 and RIPR connect the cytosolic RNA exosome to the Ski complex in *Arabidopsis*": Suppl. Table 6_Primers

Supplementary Table 6: Primers used in this study

Genotyping primers

| | |
|---|---|
| *cer7-2* fw | ATATTTGAGTGGTGCTGCTGG |
| *cer7-2* rev | AAACTCGACAAAGAGGGAAGC |
| *cer7-3* fw | AAAGCTTCCCTCTTTGTCGAG |
| *cer7-3* rev | GCCATTGGCATTAACTGTCAC |
| *rrp45a* fw | GTTGTTGGTTGCAGAGAAAGC |
| *rrp45a* rev | TGCGAGAAGTCTCAACATGTC |
| *rst1-2* fw | GCGTGTTCTAAGCCATCTTTG |
| *rst1-2* rev | GCAAGGAAATAAGAGCAAGGG |
| *rst1-3* fw | TTGATTTCATCAATGGCTTCC |
| *rst1-3* rev | CTGACAAGGGACGTTAGTTCG |
| *rst1-4* fw | TGAGGTGTCTGAAGTGGTGCA |
| *rst1-4* rev | CAAAGATGGCTTAGAACACGC, cleave product with *Ban1* |
| *riprT/C* fw | CGATGGACTCAAAATCTCTAGCTAAATCGAAGA |
| *ripT/C* rev | ACCTTGCCCGAACAACAAGA |
| *sgs3-13* fw | AAGGCCATGCTTGTACATGAG |
| *sgs3-13* rev | TATGAGGCTCTTAGAGCACGC |
| *MIM156* fw | AAGAAAAATGGCCATCCCCTAGC |
| *MIM156* rev | TGACAGAAGATAGAAGTGAGCAT |
| *gRST1* fw | GACGTGTTGATTGAGATAGT |
| *gRST1* rev | AACAGCTATGACCAT (M13 rev present in T-DNA) |

probes

| | |
|---|---|
| *CER3* fw | ACAGGTAATCTCAACTCCGAGG |
| *CER3* rev | TGGAACACCAGCTACGACAC |
| *IPS1* (MIM) fw | AAGAAAAATGGCCATCCCCTAGC |
| *IPS1* (MIM) rev | TAGAGGGAGATAAACAAAACTCGCAGT |
| U6 | GCTAATCTTCTCTGTATCGTTCCA |
| 7SL | ATATGAAGATCGGACCAGCAGGC |
